# Genome size evolution in the beetle genus *Diabrotica*

**DOI:** 10.1101/2021.09.04.458993

**Authors:** Dimpal Lata, Brad S Coates, Kimberly K O Walden, Hugh M Robertson, Nicholas J Miller

## Abstract

Diabrocite corn rootworms are one of the most economically significant pests of maize in the United States and Europe and an emerging model for insect-plant interactions. Genome sizes of several species in the genus *Diabrotica* were estimated using flow cytometry along with that of *Acalymma vittatum* as an outgroup. Genome sizes ranged between 1.56 - 1.64 gigabase pairs (Gb) and between 2.26-2.59 Gb, respectively, for the *Diabrotica* subgroups fucata and virgifera; the *Acalymma vittatum* genome size was around 1.65 Gb. This result indicated that a substantial increase in genome size occurred in the ancestor of the virgifera group. Further analysis of fucata group and virgifera group genome sequencing reads indicated that the genome size difference between the *Diabrotica* subgroups could be attributed to a higher content of transposable elements, mostly miniature inverted-transposable elements (MITEs) and gypsy-like long terminal repeat (LTR) retroelements.

## Introduction

The family Chrysomelidae is one of the largest families of phytophagous beetles (Order: Coleoptera), with nearly 40,000 species. A large number of species are important agricultural and forestry pests causing negative economic impacts (Reid 1995; Nie *et al*. 2020). A subtribe of Chrysomelidae, Diabroticina includes important agricultural pests from the genera *Acalymma, Cerotoma*, and *Diabrotica* (Toepfer *et al*. 2009). Many species of the genus *Acalymma* are specialists on Cucurbitaceae, with *Acalymma vittatum*, the striped cucumber beetle, being one of the key pests of cucurbits in the northeastern United States (Lewis *et al*. 1990). The bean leaf beetle, *Cerotoma trifurcata*, is an important pest of leguminous crops such as peas and soybeans throughout the eastern USA (Koch *et al*. 2004). *Diabrotica*, the most diverse genus (Eben and Espinosa de los Monteros 2013), includes some of the most destructive insect pests impacting US agriculture. *Diabrotica spp*. are divided into three groups: signifera, fucata, and virgifera, with the latter two containing recognized pest species (Krysan 1986). The species in the fucata group are multivoltine and polyphagous, while species in the virgifera group are univoltine and oligophagous (Branson and Krysan 1981; Krysan 1982). *Diabrotica undecimpunctata* (southern corn rootworm) within the fucata group is a generalist feeder that feeds on several crops, including cucurbits, peanuts, and maize in the southern USA (Jackson *et al*. 2005). *Diabrotica virgifera virgifera* (western corn rootworm), *Diabrotica barberi* (northern corn rootworm), and *Diabrotica virgifera zeae* (Mexican corn rootworm) from the virgifera group are specialist feeders and are pests of maize. *Diabrotica virgifera virgifera* is most abundant in the US Corn Belt but is found throughout much of the United States as well as parts of Canada and Mexico. *Diabrotica virgifera virgifera* and *Diabrotica barberi* are sympatric in the northern part of the US Corn Belt, while *D. v. zeae* is sympatric with *D. v. virgifera* over part of their range in Texas, Arizona, and Mexico (Bragard *et al*. 2019).

As the name “corn rootworm” suggests, several *Diabrotica* species cause substantial economic damage to maize agriculture. The western corn rootworm, *Diabrotica virgifera virgifera*, is considered one of the most destructive pests of maize throughout the US Corn Belt in the United States, and it accounts every year for over $1 billion in yield losses and pest management costs (Sappington *et al*. 2006). The species has also been introduced into Europe and has become widespread because of a combination of transatlantic introductions and intra-continental movement (Ciosi *et al*. 2008, 2011; Miller *et al*. 2010). *Diabrotica barberi* is a serious maize pest but is less widespread than *D. v. virgifera* (Capinera 2008). Maize-specialist corn rootworms have proven to be highly adaptable to a variety of pest management tactics. Resistance has evolved to a variety of synthetic insecticides (Ball and Weekman 1962; Meinke *et al*. 1998; Pereira *et al*. 2015; Souza *et al*. 2019, 2020) to most rootworm active transgenic maize varieties (Gassmann *et al*. 2014; Calles-Torrez *et al*. 2019) and to cultural control methods (Krysan *et al*. 1984; Gray *et al*. 2009). Although *D.undecimpunctata* is more widely distributed, it is unable to survive the winter temperatures of the US Corn Belt, and it is considered an occasional pest of maize.

The genus *Diabrotica* is emerging as a model for insect-plant interactions in generalist *versus* specialist herbivory. The ancestral state for the genus is thought to be generalist feeding with a host plant range that includes Cucurbitaceae, Fabaceae, and Poaceae, which is retained in the fucata group (Eben and Espinosa de los Monteros 2013). Following the split between the fucata and virgifera groups, around 30 million years ago, the virgifera group specialized on Poaceae (Eben and Espinosa de los Monteros 2013). Consequently, pest *Diabrotica* includes both generalist and specialist species that share a common, experimentally-tractable host plant in maize.

The benzoxazinoid DIMBOA (2,4-dihydroxy-7-methoxy-1,4-benzoxazin-3-one) is one of the major secondary metabolites produced by maize plants (Sasai *et al*. 2009). Specialist virgifera group species and generalist fucata group species respond distinctly to this metabolite. *Diabrotica virgifera virgifera* larvae gained significantly more dry weight when fed wild-type plants compared to larvae fed mutant plants, deficient for

DIMBOA biosynthesis. However, *D. undecimpunctata* performed equally well when fed on both types of plants (Alouw and Miller 2015). The enhanced performance of specialist *D. v. virgifera* may be related to its ability to use DIMBOA as a signal to locate nutritious parts of roots, while the generalist from the fucata group does not (Robert *et al*. 2012). Further, RNA-Seq studies showed transcripts encoding for a CYP9-like cytochrome P450 monooxygenase were expressed in *D. v. virgifera* larvae feeding on wild type plants but not in larvae feeding on benzoxazinoid-deficient mutant plants (Miller and Zhao 2015), suggesting a cytochrome P450 mediated adaptation to benzoxazinoids in *D. v. virgifera*.

Given the economic importance of and growing research interest in *Diabrotica* beetles, there has been considerable interest in obtaining sequences of their genomes and understanding genetic mechanisms for adaptations. Most *Diabrotica* genetics and genomics research so far has been concentrated on *D. v. virgifera* (Gray *et al*. 2009; Miller *et al*. 2010). The genome of *D. v. virgifera* is one of the larger genomes among beetles and is estimated to be around 2.58 Gb (Coates *et al*. 2012), whereas the average genome size for Coleoptera is 0.76Gb (Schoville *et al*. 2018; Gregory 2021). Increased sizes of eukaryotic genomes are generally attributed to corresponding increased numbers of repetitive DNA elements (Kidwell 2002), where a large proportion of repeats are composed of different transposable element (TE) sequences (Kojima 2019). Eukaryotic transposons are divided into retroelements that propagate by an RNA intermediate (class I) and DNA elements (class II) that mobilize by a “cut-and-paste” mechanism (Finnegan 1989; Wicker *et al*. 2007).

There is evidence that the *D. v. virgifera* genome contains a high proportion of repetitive elements (Coates *et al*. 2012, 2014). The cadherin gene of *D. v. virgifera* is approximately 13.3 fold larger than the *Tribolium castaneum* ortholog due to much larger introns. The presence of numerous MITE-like elements within the cadherin gene of *D. v. virgifera* indicates that the difference in the gene size is due to the insertion of transposable elements in the *D. v. virgifera* introns (Coates *et al*. 2012). Class I BEL-like long terminal repeat (LTR) retrotransposons have been also found in the *D. v. virgifera* genome (Coates *et al*. 2014). Initially, MITEs were found as key components of plant genomes, as they are frequently associated with genes with high copy numbers indicating a possible role in gene expression and genome evolution (Santiago *et al*. 2002; Oki *et al*. 2008). They are also found in animals, including mosquitoes, *Drosophila*, fish, and humans (Deprá *et al*. 2012). Similar to MITES, LTR retrotransposons were also first discovered in plants. They are usually located largely in intergenic regions and are often the single largest component of plant genomes (Kumar and Bennetzen 1999; Feschotte *et al*. 2002). Previous studies have also revealed that both MITES and LTR retrotransposons can modify gene expression by inserting into promoter regions (Butelli *et al*. 2012; Wang *et al*. 2017; Liu *et al*. 2019). TE integration and excision can introduce novel variation (McClintock 1950; Wendel and Wessler 2000). Transposons can cause mutations by inserting themselves into functional regions and causing change by either modifying or eliminating gene expression (Feschotte 2008; Oliver and Greene 2009). They may also lead to genomic rearrangement (Maumus *et al*. 2015; Mat Razali *et al*. 2019).

Although the genome size of *D. v. virgifera* has been reported, no genome size data have been obtained for the other species in the genus *Diabrotica* and related genera. Since the genome size of *D. v. virgifera* is relatively large, we hypothesized that there has been a recent expansion in genome size in the lineage leading to it. We tested this hypothesis by estimating and comparing the genome sizes of *D. v. virgifera* with those of several *Diabrotica* species and an outgroup species, *A. vittatum*. As a high proportion of repetitive elements were found in the cadherin gene of *D. v. virgifera*, we further hypothesized that genome size expansion in the lineage leading to *D.v. virgifera* was due to a general increase in repetitive elements. To test this hypothesis, we looked at the nature and quantity of repetitive elements in the virgifera group and compared it with the fucata group.

## Materials and Methods

### Sample Collection for Flow Cytometry

Specimens of *D. v. virgifera* and *A. vittatum* were collected from a maize field in Illinois in 2017, while *D. barberi* were collected from Wisconsin by Tracy Schilder, Wisconsin Department of Agriculture. Specimens of *D. v. zeae* were collected from Texas by Thomas Sappington, US Department of Agriculture Agricultural Research Service and *D. balteata* were provided by Blair Siegfried and Heather McAuslane, University of Florida, from a laboratory colony. *Diabrotica undecimpunctata* were obtained from Crop Characteristics (Farmington, Minnesota, USA). Adult male *Periplaneta americana*, which were used as an external reference (Guo *et al*. 2015; He *et al*. 2016) for flow cytometric measurement, were obtained from Carolina Biological Supply (Burlington, North Carolina, USA). All samples were flash-frozen in liquid nitrogen and preserved at - 80°C.

### Sample Preparation for Flow Cytometry

Genome size estimates were generated for eight individuals from five species of *Diabrotica* and one species of *Acalymma*. Preparations of nuclei were based on the method of Hare and Johnston (2012). The heads of single individuals were homogenized in 1ml of cold Galbraith buffer (4.26 g of MgCl_2_, 8.84 g of sodium citrate, 4.2g of MOPS, 1ml of Triton-x, and 1mg of boiled ribonuclease A into 1 liter of ddH20) placed in a 7ml Kontes Dounce. The homogenate was filtered through a 20 μm nylon mesh in a 1.5ml Eppendorf tube. Nuclei were stained with propidium iodide (PI) at 50 μg/ml in the dark at 4°C for an hour. In addition to the test sample, the brain tissue of *P. americana* was used as a standard (Hanrahan and Johnston 2011). The brain tissue of *P. americana* was dissected out, and the nuclear suspension was prepared and stained as described above.

### Flow Cytometric Analysis

Stained nuclei were analyzed using an Attune NxT Flow Cytometer (Thermo Fisher Scientific, Waltham, Massachusetts). The propidium iodide-stained nuclei were excited by exposing them to the 488 nm blue laser. Red fluorescence from the propidium iodide was collected using the YL2 detector channel. The calibration of the flow cytometer was performed using a standard manufacturer’s protocol before use. During each sample run, the linearity of the fluorescence measurement was confirmed by checking that the mean channel number of the 4C nuclei (G2 phase) was double that of 2C nuclei (G1 phase). At least 1000 nuclear events were collected under each unknown and standard 2C peak. The nuclei peak (PI fluorescence histogram) and coefficient of variation (CV) for each peak of interest (sample and standard) were obtained using the gating function in the Attune Software. The coefficient of variation (CV) was less than 5 percent which is considered appropriate for accurate genome size estimates (Dolezel *et al*. 2007; Tomaszewska *et al*. 2021). The known genome size of the external standard (3.34Gb, Harahan and Johnson 2011) and the relative fluorescence obtained from the sample and external standard were then used to estimate the genome size using the following formula:

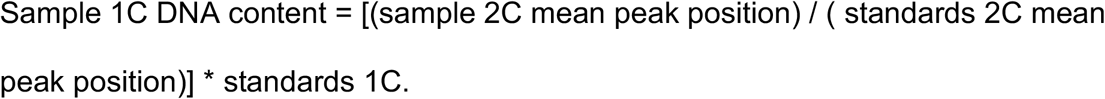

Genome size variations were analyzed using analysis of variance (ANOVA) and Tukey’s honest significant difference (HSD) post-hoc analyses using R statistical software (version: 4.10) (R Core Team 2021). Letters were assigned showing significance based on Tukey HSD post-hoc test using R statistical software (version: 4.10) (R Core Team 2021).

### Sampling and Genomic DNA Sequencing

Separate sample collection and preparation were done to obtain the data from Illumina whole-genome shotgun sequencing which were used to analyze the repetitive DNA content of *D. barberi*, *D.undecimpunctata*, and *D. v. virgifera*. Adult *D. barberi* (*n* = 71) and *D. undecimpunctata* (*n* = 50) were collected from maize fields near Ames, Iowa, and Monmouth, Illinois, respectively. Each sample was pooled by species, flash-frozen in liquid nitrogen and ground in liquid nitrogen, and then DNA extracted from ~3.0 mg of tissue using the Qiagen DNeasy Blood and Tissue Extraction kit, with modifications as described (Coates *et al*. 2014). Two micrograms of extracted DNA was submitted to the Iowa State University DNA Facility (Ames, IA, USA) from which ~500bp insert indexed sequencing libraries were generated using the Illumina TruSeq v2 Library Construction Kit (Illumina, San Diego, CA). Single-end 100-bp reads were generated from *D. barberi* and *D. undecimpunctata* libraries in separate lanes of an Illumina HiSeq2500. Raw reads were submitted to the Biotechnology Information (NCBI) Short Read Archive (SRA) under accessions SRR13363759 and SRR13364002 for *D. barberi* and *D. undecimpunctata*, respectively.

*Diabrotica virgifera virgifera* adult females of inbred line Ped12, developed by the USDA-ARS North Central Agricultural Research Laboratory were used for the genomic DNA isolation. Briefly, whole beetles were homogenized in an SDS-based cell lysis solution followed by overnight incubation with Proteinase K at 55°C. Cellular debris was pelleted and RNA was digested with RNaseA. The homogenate was mixed with a high-salt solution and incubated overnight at 4°C. The DNA in the supernatant was precipitated overnight with ethanol at −20°C. DNA was quantified on an Invitrogen Qubit. A paired-end short-insert genomic DNA library was prepared at the Roy J. Carver Biotechnology Center at the University of Illinois at Urbana-Champaign using an Illumina TruSeq DNAseq Sample Prep kit. Reads were sequenced to 100bp with the Illumina TruSeq SBS sequencing kit version 3 on an Illumina HiSeq 2000 instrument using Casava 1.8 for basecalling. Raw reads were submitted to the Biotechnology Information (NCBI) Short Read Archive (SRA) under accession SRR6985755. Sequencing generated 90 million single-end reads of 100bp for *D. barberi*, 118 million single-end reads of 100bp for *D.undecimpunctata*, and 116 million 100-bp paired-end reads for *D. v. virgifera*.

### Annotation and Quantification of Repeat Content from sequencing data

Raw reads were quality-filtered using fastp software (version 0.20.1) with a minimum 20 average Phred score. Reads mapping to mitochondrial genome sequences of *Diabrotica* species available through the NCBI website (KF658070.1, KF669870.1) were identified (minimap2 v2.17) and filtered out as implemented in the SSRG workflow (Pombert, 2021). Repetitive elements in the genomes of *D.undecimpunctata*, *D. barberi*, and *D. v. virgifera* were assembled and quantified using dnaPipeTE v1.3 (Goubert *et al*. 2015) and annotated using the DeepTE tool (Yan *et al*. 2020). To quantify the proportion of TEs, dnaPipeTE uses samples of sequence reads instead of genome assemblies, making this pipeline (dnaPipeTE) applicable for genomes with lower sequencing depth. The pipeline performed assembly of repetitive reads into contigs from low coverage sampling of raw reads using Trinity (Grabherr *et al*. 2011) and annotated them using RepeatMasker (Smit *et al*. 1996-2010) with built-in Repbase libraries (Bao *et al*. 2015, version 2017-01-27). Quantification was done by mapping a random sample of reads onto the assembled repeats. The parameters set as the benchmark for repeat content analysis for genomes greater than 500Mb (Goubert *et al*. 2015), including the coverage parameter, were used to run dnaPipeTE. The pipeline was run for all three species using 0.1x coverage. Additionally, 0.1x coverage was chosen based on the high N50 metric and plateauing point of transposable elements i.e, increasing the coverage beyond 0.1x only marginally increased the proportion of transposable elements for all three species. The dnaPipeTE pipeline does not annotate novel repeats that do not match an entry in the included Repbase library. A high proportion of repeats from each of the three beetle species were not annotated by dnaPipeTE. DeepTE, a deep learning method based on convolutional neural networks, was used to classify and annotate the unknown TEs. DeepTE uses eight trained models to classify TEs into superfamilies and orders. All the TE contigs assembled by dnaPipeTE were analyzed using DeepTE, whether or not they had been previously classified by dnaPipeTE. Combining the results of the assembly and quantification by dnaPipeTE with the classification results from DeepTE allowed the abundance of repeat families in the genomes of all three species to be determined.

For comparison with the dnaPipeTE *de-novo* assembly of repetitive elements, the percentage of repetitive elements in the *D. v. virgifera* genome assembly (NCBI RefSeq accession GCF_003013835.1) was analyzed with RepeatModeler version 2.0.1 (Flynn *et al*. 2019). Repeatmodeler is a *de-novo* transposable element identification package that uses three repeat finding programs (RECON, RepeatScout, and LtrHarvest/Ltr_retriever) to discover repetitive DNA sequences in the genome. These repetitive DNA sequences were annotated by repeatClassifier based on the similarity to RepBase and Dfam databases. The annotated library produced was used as input to RepeatMasker to detect and mask repeats in the genome. Default parameters were used to run RepeatModeler.

## Results

### Genome size

The genome sizes of *D. v. zeae, D. v. virgifera*, and *D. barberi* from the virgifera group were estimated to be 2.59Gb ± 0.01, 2.58 Gb ± 0.02, and 2.26Gb ± 0.04, respectively. The genomes of *D. balteata* and *D.undecimpunctata* from the fucata group were estimated at 1.64Gb ± 0.01 and 1.56 Gb ± 0.00. The outgroup species *A. vittatum* genome size was estimated to be 1.65 Gb ± 0.01. An analysis of variance (ANOVA) showed a significant difference (*F*_*(5, 42)*_ = 597, *p* < 0.001) in the genome sizes of the species under study. A subsequent Tukey HSD test showed that there were no significant differences in genome size between *D. v. virgifera* and *D. v. zeae*, between *D. balteata* and *D.undecimpunctata*, or between *D. balteata* and *A. vittatum*. The estimated genome size for each species with their phylogenetic relationships is shown in Figure 1.

**Figure 1:**
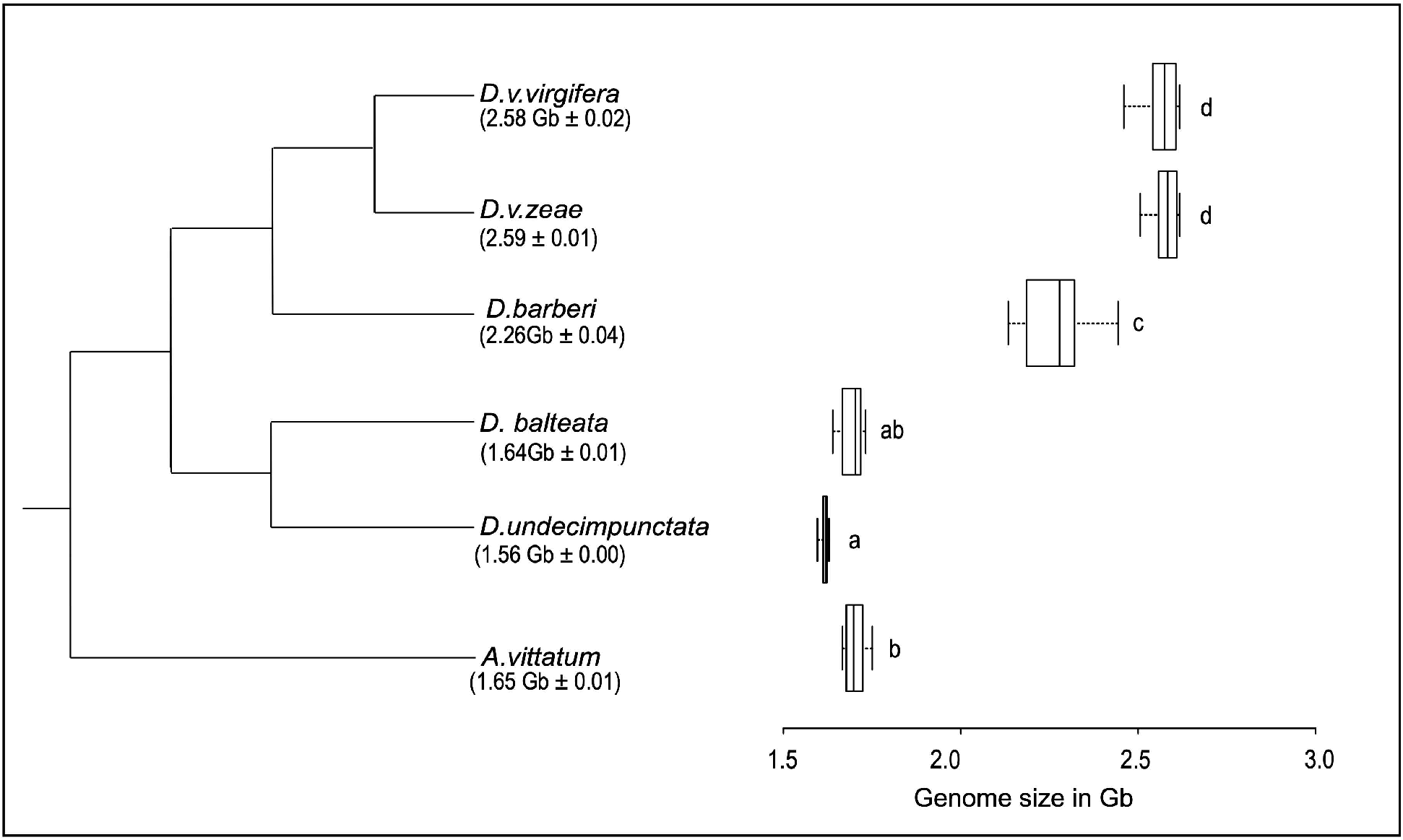
Genome size evolution within the genus *Diabrotica*. Phylogeny of *Diabrotica* and outgroup *Acalymma vittatum* is based on (Eben and Espinosa de los Monteros 2013). Letters a - d indicate groups with no significant difference in mean 1C-value (Tukey HSD; *α* = 0.05).

### Repeat Content analysis

The repeatomes of *D.v.virgifera*, *D. barberi*, and *D.undecimpunctata* comprised 72.4%, 70.3%, and 52.7% of their genomes, respectively. The repeat content obtained via Repeat Modeler for the draft *D. v. virgifera* genome was 57.4%. As the assembly of the *D.v. virgifera* genome was based on short-reads, TEs were expected to be under-represented because of the difficulty of assembling individual copies. TE-rich large genomes are difficult to assemble and often end up with high levels of fragmentation around repetitive regions leading to underestimation of TE content (Green 2002).

To further investigate the classes of repeat families that contributed to the genome size variation between the two groups of *Diabrotica*, the DeepTE annotations were coupled with the dnaPipeTE abundance quantification to estimate the abundance of different repeat elements in the genomes of *D. v. virgifera*, *D.barberi*, and *D.undecimpunctata*. The TEs that accounted for most of the difference in the genome size of the two groups were annotated as class II DNA Tc1-mariner Miniature-repeat Transposable Elements (MITEs) (ClassII_DNA_TcMar_MITE) and class II DNA hAT MITE (ClassII_DNA_hAT_MITE) of transposable elements and class I long terminal repeat (LTR) Gypsy (ClassI_LTR_Gypsy) transposable elements and is shown in Figure 2. The *D. v. virgifera* genome contained a large amount of ClassII_DNA_TcMar_MITE(0.61 Gb), ClassII_DNA_hAT_MITE (0.23 Gb), and ClassI_LTR_Gypsy(0.31 Gb) (Figure 2). Similarly, *D. barberi* also had a high amount of ClassII_DNA_TcMar_MITE(0.50 Gb), ClassII_DNA_hAT_MITE (0.19 Gb), and ClassI_LTR_Gypsy(0.25 Gb) (Figure 2). The *D. undecimpunctata* had a lower amount of ClassII_DNA_TcMar_MITE(0.15 Gb), ClassII_DNA_hAT_MITE (0.06 Gb), and ClassI_LTR_Gypsy(0.19 Gb) (Figure 2). Transposable elements from other repeat families such as nLTRS, helitrons, and others from class I and class II DNA elements were not as prominent as those mentioned above. Single-low copy sequences representing the non-repetitive portion of the genomes of *D. v. virgifera*, *D.barberi*, and *D.undecimpunctata* totaled 0.73 Gb, 0.68 Gb, and 0.75 Gb, respectively (Figure 2).

**Figure 2:**
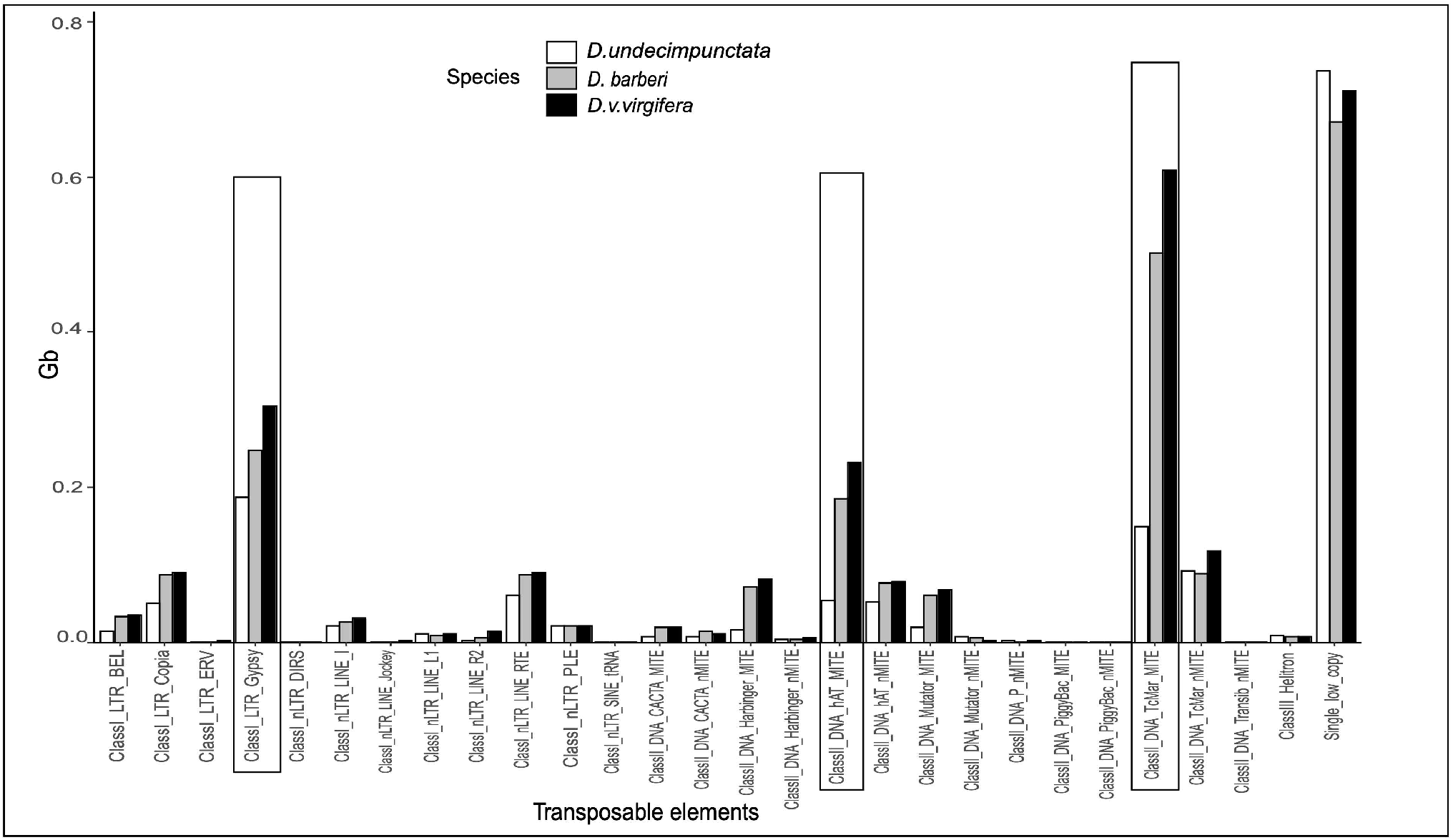
Predicted transposable elements (In Gb) in the genome of three species of *Diabrotica*. The boxed transposable elements are the top three highest contributors to the genome size variation between the groups of fucata and virgifera.

## Discussion

*Diabrotica* is the most diverse genus within the subtribe Diabroticina and includes 354 species native to America. Only species from the fucata and virgifera groups occur in the United States (Krysan 1986), while species from the signifera group are endemic to South America. The signifera group species are not of economic importance, and their biology is also mostly unknown (Clark *et al*. 2001) and so the group is understudied. The economically significant pest species in this genus either belong to the virgifera group or to the fucata group, justifying the need to study them comprehensively. The expansion and adaptability of the pest species generates a sense of urgency to study them.

Our results demonstrated that the genome size of virgifera group species are approximately 1Gb larger than that of the fucata group and *Acalymma* species. The genome size for *D. v. virgifera* obtained in this study and in a previous study (Coates *et al*. 2012) are consistent. Our genome size results, when coupled with the species’ phylogenetic relationship, indicated that an expansion in genome size occurred in the common ancestor of the virgifera group leading to *D. barberi, D. v. virgifera*, and *D. v. zeae*. There was also a significant difference in the genome sizes of *D. barberi* and *D. virgifera subsp*. suggesting a possible further expansion of the genome size in the common ancestor of *D. v. virgifera* and *D. v. zeae*.

Repeat elements of species from fucata and virgifera were studied to understand the basis of genome size expansion in *Diabrotica*. Differences in the total amount of repetitive DNA accounted almost entirely for the differences in genome size between the three species that were studied. The total amounts of low copy number DNA (which includes most genes) were very similar, only differing by a few 10s of Mb. These results strongly support the hypothesis of TE proliferation as the driver of genome expansion in the virgifera group. Our data do not support a role for genome duplication, as this mechanism would also produce substantial differences in the quantity of low copy number DNA.

The genomes of the virgifera and fucata group of *Diabrotica* species contained many common TEs, but the abundance of three TE families differed substantially and accounted for approximately 74% of the difference in genome size between the two groups and also 66% of the difference within the virgifera group between *D.barberi* and *D. virgifera subsp*. The MITE-like Tc1-mariner and hAT elements and LTR Gypsy retroelements were more abundant in the virgifera group (45% in *D. virgifera*, 42% in *D.barberi*). Miniature-repeat Transposable Elements (MITEs) are short AT-rich (<0.5kb) derivatives of DNA elements whose internal sequence lacks an open reading frame (Lu *et al*. 2012), contain conserved terminal inverted repeats, are flanked by target site duplications, and are closely associated with euchromatic genes (Kuang *et al*. 2009). Gypsy elements are one of the most abundant classes of the long terminal repeat (LTR)-retrotransposons superfamily with large numbers of copies found in almost all the plants, animals, and fungi tested (Thomas-Bulle *et al*. 2018). Generally, class I transposable elements have been reported to be in high abundance in insect genomes, such as *Tribolium castaneum* (Wang *et al*. 2008), *Drosophila* (Clark *et al*. 2007) and *Bombyx mori* (Osanai-Futahashi *et al*. 2008) in comparison to class II elements. However, in *Diabrotica*, we discovered an abundance of class II elements and, to some extent, class I elements.

Genome size varies enormously among eukaryote species (Hidalgo *et al*. 2017), including beetles in the family Chrysomelidae (Petitpierre *et al*. 1993; Hanrahan and Johnston 2011). Some have speculated that variation in genome size *per se* is related to variation in phenotypic traits in insects. Correlations have been reported between genome size and traits, including body size (Ferrari and Rai 1989; Finston *et al*. 1995; Palmer and Petitpierre 1996; Palmer *et al*. 2003), development rate (Carreras *et al*. 1991; Gregory *et al*. 2003), and indeed, host plant range (Matsubayashi and Ohshima 2015; Calatayud *et al*. 2016; Zhang *et al*. 2019). However, in many of these examples, different studies have produced contradictory results, with both positive and negative correlations for the same trait. In the case of our data, there was no relationship between host plant range and genome size. We found the cucurbit-specialist *Acalymma* had a similar-sized genome to generalists in the fucata group of *Diabrotica*, whereas the other specialist species in the study (virgifera group *Diabrotica*) had substantially larger genomes. The data from our study support the view that correlations between genome size and phenotypic traits are generally coincidental.

Our data showed that the increased genome size in the virgifera group of *Diabrotica* species was the result of the proliferation of a few TE families. Comparative genomic studies in insects have revealed that repeat elements can make large contributions to genome size variation. Variation in TE abundance can be seen both within and among species (Lynch 2007). Honeybees, with a genome size of 230Mb, show very few repeat elements, representing a case of TE extinction (Weinstock *et al*. 2006). Similarly, the small genome of *Belgica antarctica*, the Antarctic midge, (99 Mb) is also due to the reduction of repeats in the genome (Kelley *et al*. 2014). There are also cases of TE proliferations, such as in *Locusta migratoria*, that led to its large genome size of 6.5 Gb (Wang *et al*. 2014). Cases of increase in genome size as a consequence of transposable elements have also been reported in wood white (Leptidea) butterflies and North American fireflies (Lampyridae) (Lower *et al*. 2017; Talla *et al*. 2017). Overall, it appears that genome size in insects is fairly plastic and largely driven by the loss and gain of transposable elements.

It is likely that the proliferation of TEs responsible for the increase in genome size observed in our study occurred sometime between the divergence of the ancestors of the virgifera and fucata groups 30 million years ago, but before the radiation of the virgifera group species, around 17 million years ago (Eben and Espinosa de los Monteros 2013). Host genomes have several mechanisms to suppress transposable element expression and mobility (Bourque *et al*. 2018), including epigenetic silencing through histone modifications or DNA methylation, targeted mutagenesis, small RNA interference, as well as sequence-specific repressors such as the recently profiled KRAB zinc-finger proteins (Fouché *et al*. 2020; Maupetit-Mehouas and Vaury 2020). At the same time, some transposable elements have evolved regulatory sequences controlling their own copy number to autonomously replicate in the genome (Lohe and Hartl 1996; Saha *et al*. 2015). TE de-repression is triggered by environmental stimuli, in particular stress (Bundo *et al*. 2014; Voronova *et al*. 2014; Fouché *et al*. 2020), impacting transcription levels and increasing transpositional activity (Dubin *et al*. 2018). Additionally, there are other factors influencing TE mobilization, for example, demethylation and the removal of repressive histone marks during epigenetic reprogramming stages (Russell and LaMarre 2018). The factors that led to de-repression and subsequent proliferation of TEs in the ancestor of the virgifera group of species are unknown.

Although gross variation in genome size, driven by gain and loss of TEs, does not correlate with phenotypic adaptation, mutations associated with specific TE insertions or excisions can be adaptive. Mostly, TE insertions are presumed to be deleterious or neutral, but some have been shown to be selectively advantageous. There are several studies showing that TE-mediated insertions have led to insecticide resistance. In pink bollworm *Pectinophora gossypiella*, a major pest of cotton (Rostant *et al*. 2012), several independent TE insertions in the *PgCad1* gene conferred resistance to *Bt* Cry1Ac toxin (Fabrick *et al*. 2011; Wang *et al*. 2019). Cases of resistance to *Bacillus thuringiensis* (Bt) toxins have also been reported in *Heliothis virescens* which is caused by disruption of a cadherin-superfamily gene by TE insertion (Gahan 2001). TE insertions in xenobiotic metabolism related genes such as those encoding cytochrome P450 monooxygenases and glutathione S-transferases in *Helicoverpa armigera* are the cause of resistance to insecticides (Klai *et al*. 2020). Another example demonstrating that TEs can produce adaptive mutations has been reported in *D. melanogaster* (Rostant *et al*. 2012; Gilbert *et al*. 2021). An increased resistance to dichlorodiphenyltrichloroethane (DDT) in *D. melanogaster* has been reported due to Cyp6g1 upregulation caused by insertion of the Accord transposon in the 5’ regulatory region of the Cyp6g1 gene (Chung *et al*. 2007). Similarly, TE insertion, which truncates the CHKov1 gene in *D. melanogaster*, confers resistance towards organophosphate (Aminetzach 2005).

The degree to which mutations caused by the proliferation of TEs in the virgifera group of *Diabrotica* contributed directly to the evolution of the group is unknown. Tackling this question will require a comparative evolutionary genomic analysis of the *Diabrotica* genus. To date, significant genome sequence data are only available for one species, *Diabrotica virgifera virgifera*. However, the USDA-ARS Ag100Pest initiative aims to sequence additional *Diabrotica* genomes. The information on the evolution of size and repeat content of *Diabrotica* genomes presented in this paper will help to inform the optimum strategies for sequencing additional genomes for the genus and, potentially, Diabroticite genomes more generally.

## Data availability

Nucleic acid sequencing data are available from the NCBI Sequence Read Archive under the accession numbers provided above. All flow cytometry data are available for download from Figshare.

## Acknowledgments

We would like to thank Dr. Thomas Sappington, Dr. Blair Siegfried, Dr. Heather McAuslane, and Tracy Schilder for providing insect samples. We thank Dr. J. Spencer Johnston and David Leclerc for their advice and guidance on flow cytometry methods and Dr. Clément Goubert’s advice on dnaPipeTE. Additionally, we also thank two anonymous reviewers for their constructive feedback on an earlier version of the manuscript.

## Funding and Competing Interests

This work was supported and funded by USDA NIFA award number 2011-67009-30134 and by the USDA Agricultural Research Service (ARS) (CRIS Project 5030-22000-019-00D). This article reports the results of research only, and mention of any commercial product or service does not constitute a recommendation for its use by USDA. The findings and conclusions in this publication are those of the authors and should not be construed to represent any official USDA or United States government determination of policy. USDA is an equal opportunity employer and provider.

## Notes

### Competing Interest Statement

The authors have declared no competing interest.

### Summary of Updates

Updated in response to 2 anonymous reviewers' comments

## References

Alouw, J. C., and N. J. Miller, 2015 Effects of benzoxazinoids on specialist and generalist *Diabrotica* species. J. Appl. Entomol. 139: 424–431.

Aminetzach, Y. T., 2005 Pesticide resistance via transposition-mediated adaptive gene truncation in *Drosophila*. Science 309: 764–767.

Ball, H. J., and G. T. Weekman, 1962 Insecticide resistance in the adult western corn rootworm in Nebraska. J. Econ. Entomol. 55: 439–441.

Bao, W., K. K. Kojima, and O. Kohany, 2015 Repbase Update, a database of repetitive elements in eukaryotic genomes. Mob. DNA 6: 11.

Bourque, G., K. H. Burns, M. Gehring, V. Gorbunova, A. Seluanov et al., 2018 Ten things you should know about transposable elements. Genome Biol. 19: 199.

Bragard, C., K. Dehnen-Schmutz, F. Di Serio, P. Gonthier, M. Jacques et al., 2019 Pest categorisation of *Diabrotica virgifera zeae*. EFSA J. 17:.

Branson, T. F., and J. L. Krysan, 1981 Feeding and oviposition behavior and life cycle strategies of *Diabrotica*: an evolutionary view with implications for pest management. Environ. Entomol. 10: 826–831.

Bundo, M., M. Toyoshima, Y. Okada, W. Akamatsu, J. Ueda et al., 2014 Increased l1 retrotransposition in the neuronal genome in schizophrenia. Neuron 81: 306–313.

Butelli, E., C. Licciardello, Y. Zhang, J. Liu, S. Mackay et al., 2012 Retrotransposons control fruit-specific, cold-dependent accumulation of anthocyanins in blood oranges. Plant Cell 24: 1242–1255.

Calatayud, P.-A., C. Petit, N. Burlet, S. Dupas, N. Glaser et al., 2016 Is genome size of Lepidoptera linked to host plant range? Entomol. Exp. Appl. 159: 354–361.

Calles-Torrez, V., J. J. Knodel, M. A. Boetel, B. W. French, B. W. Fuller et al., 2019 Field-evolved resistance of northern and western corn rootworm (Coleoptera: Chrysomelidae) populations to corn hybrids expressing single and pyramided Cry3Bb1 and Cry34/35Ab1 Bt proteins in North Dakota. J. Econ. Entomol. 112: 1875–1886.

Capinera, J. L., 2008 Northern corn rootworm, *Diabrotica barberi* Smith & Lawrence (Coleoptera: Chrysomelidae), pp. 2616–2619 in Encyclopedia of Entomology, edited by J. L. Capinera. Springer Netherlands, Dordrecht.

Carreras, I., A.-F. A., C. Juan, and E. Petitpierre, 1991 Tasa de desarrollo en *Tribolium brevicornis* (Coleoptera: Tenebrionidae) y su relación con el tamaño del genoma.

Chung, H., M. R. Bogwitz, C. McCart, A. Andrianopoulos, R. H. Ffrench-Constant et al., 2007 *Cis*-regulatory elements in the *Accord* retrotransposon result in tissue specific expression of the *Drosophila melanogaster* insecticide resistance gene *Cyp6g1*. Genetics 175: 1071–1077.

Ciosi, M., N. J. Miller, K. S. Kim, R. Giordano, A. Estoup et al., 2008 Invasion of Europe by the western corn rootworm, *Diabrotica virgifera virgifera:* multiple transatlantic introductions with various reductions of genetic diversity. Mol. Ecol. 17: 3614–3627.

Ciosi, M., N. J. Miller, S. Toepfer, A. Estoup, and T. Guillemaud, 2011 Stratified dispersal and increasing genetic variation during the invasion of Central Europe by the western corn rootworm, *Diabrotica virgifera virgifera*. Evol. Appl. 4: 54–70.

Clark, A. G., M. B. Eisen, D. R. Smith, C. M. Bergman, B. Oliver et al., 2007 Evolution of genes and genomes on the *Drosophila* phylogeny. Nature 450: 203–218.

Clark, T. L., L. J. Meinke, and J. E. Foster, 2001 Molecular phylogeny of *Diabrotica* beetles (Coleoptera: Chrysomelidae) inferred from analysis of combined mitochondrial and nuclear DNA sequences. Insect Mol. Biol. 10: 303–314.

Coates, B. S., A. P. Alves, H. Wang, K. K. O. Walden, B. W. French et al., 2012 Distribution of genes and repetitive elements in the *Diabrotica virgifera virgifera* genome estimated using BAC sequencing. J. Biomed. Biotechnol. 2012: 1–9.

Coates, B. S., L. M. Fraser, B. W. French, and T. W. Sappington, 2014 Proliferation and copy number variation of BEL-like long terminal repeat retrotransposons within the *Diabrotica virgifera virgifera* genome. Gene 534: 362–370.

Deprá, M., A. Ludwig, V. L. Valente, and E. L. Loreto, 2012 Mar, a MITE family of hAT transposons in *Drosophila*. Mob. DNA 3: 13.

Dolezel, J., J. Greilhuber, and J. Suda, 2007 Flow Cytometry with Plant Cells: Analysis of Genes, Chromosomes and Genomes.

Dubin, M. J., O. Mittelsten Scheid, and C. Becker, 2018 Transposons: a blessing curse. Curr. Opin. Plant Biol. 42: 23–29.

Eben, A., and A. Espinosa de los Monteros, 2013 Tempo and mode of evolutionary radiation in Diabroticina beetles (genera *Acalymma*, *Cerotoma*, and *Diabrotica*). ZooKeys 207–321.

Fabrick, J. A., L. G. Mathew, B. E. Tabashnik, and X. Li, 2011 Insertion of an intact CR1 retrotransposon in a cadherin gene linked with Bt resistance in the pink bollworm, *Pectinophora gossypiella*. Insect Mol. Biol. 20: 651–665.

Ferrari, J. A., and K. S. Rai, 1989 Phenotypic correlates of genome size variation in *Aedes albopictus*. Evol. Int. J. Org. Evol. 43: 895–899.

Feschotte, C., 2008 Transposable elements and the evolution of regulatory networks. Nat. Rev. Genet. 9: 397–405.

Feschotte, C., N. Jiang, and S. R. Wessler, 2002 Plant transposable elements: where genetics meets genomics. Nat. Rev. Genet. 3: 329–341.

Finnegan, D. J., 1989 Eukaryotic transposable elements and genome evolution. Trends Genet. TIG 5: 103–107.

Finston, T. L., P. D. N. Hebert, and R. B. Foottit, 1995 Genome size variation in aphids. Insect Biochem. Mol. Biol. 25: 189–196.

Flynn, J. M., R. Hubley, C. Goubert, J. Rosen, A. G. Clark et al., 2019 RepeatModeler2: automated genomic discovery of transposable element families: Genomics preprint.

Fouché, S., T. Badet, U. Oggenfuss, C. Plissonneau, C. S. Francisco et al., 2020 Stress-Driven Transposable Element De-repression Dynamics and Virulence Evolution in a Fungal Pathogen. Mol. Biol. Evol. 37: 221–239.

Gahan, L. J., 2001 Identification of a gene associated with Bt resistance in *Heliothis virescens*. Science 293: 857–860.

Gassmann, A. J., J. L. Petzold-Maxwell, E. H. Clifton, M. W. Dunbar, A. M. Hoffmann et al., 2014 Field-evolved resistance by western corn rootworm to multiple *Bacillus thuringiensis* toxins in transgenic maize. Proc. Natl. Acad. Sci. 111: 5141–5146.

Gilbert, C., J. Peccoud, and R. Cordaux, 2021 Transposable elements and the evolution of insects. Annu. Rev. Entomol. 66: 355–372.

Goubert, C., L. Modolo, C. Vieira, C. ValienteMoro, P. Mavingui et al., 2015 De novo assembly and annotation of the Asian tiger mosquito (*Aedes albopictus*) repeatome with dnaPipeTE from raw genomic reads and comparative analysis with the yellow fever mosquito (*Aedes aegypti*). Genome Biol. Evol. 7: 1192–1205.

Grabherr, M. G., B. J. Haas, M. Yassour, J. Z. Levin, D. A. Thompson et al., 2011 Full-length transcriptome assembly from RNA-seq data without a reference genome. Nat. Biotechnol. 29: 644–652.

Gray, M. E., T. W. Sappington, N. J. Miller, J. Moeser, and M. O. Bohn, 2009 Adaptation and invasiveness of western corn rootworm: intensifying research on a worsening pest. Annu. Rev. Entomol. 54: 303–321.

Green, P., 2002 Whole-genome disassembly. Proc. Natl. Acad. Sci. U. S. A. 99: 4143–4144.

Gregory, T. R., 2021 Animal genome size database.

Gregory, T. R., O. Nedvěd, and S. Adamowicz, 2003 C-value estimates for 31 species of ladybird beetles (Coleoptera: Coccinellidae). Hereditas 139: 121–7.

Guo, L. T., S. L. Wang, Q. J. Wu, X. G. Zhou, W. Xie et al., 2015 Flow cytometry and K-mer analysis estimates of the genome sizes of *Bemisia tabaci* B and Q (Hemiptera: Aleyrodidae). Front. Physiol. 6:.

Hanrahan, S. J., and J. S. Johnston, 2011 New genome size estimates of 134 species of arthropods. Chromosome Res. 19: 809–823.

Hare, E. E., and J. S. Johnston, 2012 Genome size determination using flow cytometry of propidium iodide-stained nuclei, pp. 3–12 in Molecular Methods for Evolutionary Genetics, edited by V. Orgogozo and M. V. Rockman. Methods in Molecular Biology, Humana Press, Totowa, NJ.

He, K., K. Lin, G. Wang, and F. Li, 2016 Genome sizes of nine insect species determined by flow cytometry and k-mer analysis. Front. Physiol. 7:.

Hidalgo, O., J. Pellicer, M. Christenhusz, H. Schneider, A. Leitch et al., 2017 Is there an upper limit to genome size? Trends Plant Sci. 22:.

Jackson, D. M., K. A. Sorensen, C. E. Sorenson, and R. N. Story, 2005 Monitoring cucumber beetles in sweetpotato and cucurbits with kairomone-baited traps. J. Econ. Entomol. 98: 12.

Kelley, J. L., J. T. Peyton, A.-S. Fiston-Lavier, N. M. Teets, M.-C. Yee et al., 2014 Compact genome of the Antarctic midge is likely an adaptation to an extreme environment. Nat. Commun. 5: 4611.

Kidwell, M. G., 2002 Transposable elements and the evolution of genome size in eukaryotes. Genetica 115: 49–63.

Klai, K., B. Chénais, M. Zidi, S. Djebbi, A. Caruso et al., 2020 Screening of *Helicoverpa armigera* mobilome revealed transposable element insertions in insecticide resistance genes. Insects 11: 879.

Koch, R. L., E. C. Burkness, and W. D. Hutchison, 2004 Confirmation of bean leaf beetle, *Cerotoma trifurcata*, feeding on cucurbits. J. Insect Sci. 4: 5.

Kojima, K. K., 2019 Structural and sequence diversity of eukaryotic transposable elements. Genes Genet. Syst. 94: 233–252.

Krysan, J. L., 1982 Diapause in the nearctic species of the *virgifera* group of *Diabrotica:* evidence for tropical origin and temperate adaptations. Ann. Entomol. Soc. Am. 75: 136–142.

Krysan, J. L., 1986 Introduction: biology, distribution, and identification of pest *Diabrotica*, pp. 1–23 in Methods for the Study of Pest Diabrotica, edited by J. L. Krysan and T. A. Miller. Springer Series in Experimental Entomology, Springer, New York, NY.

Krysan, J. L., J. J. Jackson, and A. C. Lew, 1984 Field termination of egg diapause in *Diabrotica* with new evidence of extended diapause in *D. barberi* (Coleoptera: Chrysomelidae). Environ. Entomol. 13: 1237–1240.

Kuang, H., C. Padmanabhan, F. Li, A. Kamei, P. B. Bhaskar et al., 2009 Identification of miniature inverted–repeat transposable elements (MITEs) and biogenesis of their siRNAs in the Solanaceae: new functional implications for MITEs. Genome Res. 19: 42–56.

Kumar, A., and J. L. Bennetzen, 1999 Plant retrotransposons. Annu. Rev. Genet. 33: 479–532.

Lewis, P. A., R. L. Lampman, and R. L. Metcalf, 1990 Kairomonal attractants for *Acalymma vittatum* (Coleoptera: Chrysomelidae). Environ. Entomol. 19: 8–14.

Liu, Y., M. Tahir ul Qamar, J.-W. Feng, Y. Ding, S. Wang et al., 2019 Comparative analysis of miniature inverted-repeat transposable elements (MITEs) and long terminal repeat (LTR) retrotransposons in six *Citrus* species. BMC Plant Biol. 19: 140.

Lohe, A. R., and D. L. Hartl, 1996 Autoregulation of mariner transposase activity by overproduction and dominant-negative complementation. Mol. Biol. Evol. 13: 549–555.

Lower, S. S., J. S. Johnston, K. F. Stanger-Hall, C. E. Hjelmen, S. J. Hanrahan et al., 2017 Genome size in North American fireflies: substantial variation likely driven by neutral processes. Genome Biol. Evol. 9: 1499–1512.

Lu, C., J. Chen, Y. Zhang, Q. Hu, W. Su et al., 2012 Miniature inverted-repeat transposable elements (MITEs) have been accumulated through amplification bursts and play important roles in gene expression and species diversity in *Oryza sativa*. Mol. Biol. Evol. 29: 1005–1017.

Lynch, M., 2007 The origins of genome architecture. Sinauer 7.

Mat Razali, N., B. H. Cheah, and K. Nadarajah, 2019 Transposable elements adaptive role in genome plasticity, pathogenicity and evolution in fungal phytopathogens. Int. J. Mol. Sci. 20: 3597.

Matsubayashi, K. W., and I. Ohshima, 2015 Genome size increase in the phytophagous ladybird beetle *Henosepilachna vigintioctomaculata* species complex (Coleoptera: Coccinellidae): genome size increase in ladybird beetle. Entomol. Sci. 18: 134–137.

Maumus, F., A.-S. Fiston-Lavier, and H. Quesneville, 2015 Impact of transposable elements on insect genomes and biology. Curr. Opin. Insect Sci. 7: 30–36.

Maupetit-Mehouas, S., and C. Vaury, 2020 Transposon Reactivation in the Germline May Be Useful for Both Transposons and Their Host Genomes. Cells 9: 1172.

McClintock, B., 1950 The origin and behavior of mutable loci in maize. Proc. Natl. Acad. Sci. U. S. A. 36: 344–355.

Meinke, L. J., B. D. Siegfried, R. J. Wright, and L. D. Chandler, 1998 Adult susceptibility of Nebraska western corn rootworm (Coleoptera: Chrysomelidae) populations to selected insecticides. J. Econ. Entomol. 91: 594–600.

Miller, N. J., S. Richards, and T. W. Sappington, 2010 The prospects for sequencing the western corn rootworm genome. J. Appl. Entomol. 134: 420–428.

Miller, N. J., and Z. Zhao, 2015 Transcriptional responses of *Diabrotica virgifera virgifera* larvae to benzoxazinoids. J. Appl. Entomol. 139: 416–423.

Nie, R., C. Andújar, C. Gómez-Rodríguez, M. Bai, H.-J. Xue et al., 2020 The phylogeny of leaf beetles (Chrysomelidae) inferred from mitochondrial genomes. Syst. Entomol. 45: 188–204.

Oki, N., K. Yano, Y. Okumoto, T. Tsukiyama, M. Teraishi et al., 2008 A genome-wide view of miniature inverted-repeat transposable elements (MITEs) in rice, *Oryza sativa ssp.japonica*. Genes Genet. Syst. 83: 321–329.

Oliver, K. R., and W. K. Greene, 2009 Transposable elements: powerful facilitators of evolution. BioEssays News Rev. Mol. Cell. Dev. Biol. 31: 703–714.

Osanai-Futahashi, M., Y. Suetsugu, K. Mita, and H. Fujiwara, 2008 Genome-wide screening and characterization of transposable elements and their distribution analysis in the silkworm, *Bombyx mori*. Insect Biochem. Mol. Biol. 38: 1046–1057.

Palmer, M., and E. Petitpierre, 1996 Relationship of genome size to body size in *Phylan semicostatus* (Coleoptera: tenebrionidae). Ann. Entomol. Soc. Am. 89: 221–225.

Palmer, M., E. Petitpierre, and J. Pons, 2003 Test of the correlation between body size and DNA content in *Pimelia* (Coleoptera: Tenebrionidae) from the Canary islands. Eur. J. Entomol. 100: 123–129.

Pereira, A. E., H. Wang, S. N. Zukoff, L. J. Meinke, B. W. French et al., 2015 Evidence of field-evolved resistance to bifenthrin in western corn rootworm (*Diabrotica virgifera virgifera* LeConte) populations in western Nebraska and Kansas. PLOS ONE 10: e0142299.

Petitpierre, E., C. Segarra, and C. Juan, 1993 Genome Size and Chromosomal Evolution in Leaf Beetles (Coleoptera, Chrysomelidae). Hereditas 119: 1–6.

Pombert, J.-F., 2021 PombertLab/SSRG. Pombert Lab. https://github.com/PombertLab/SSRG (Original work published 2016)

R Core Team, 2021 R: A language and environment for statistical computing. R Foundation for Statistical Computing, Vienna, Austria.

Reid, C. A. M., 1995 A cladistic analysis of subfamilial relationships I in the Chrysomelidae sensu lato (Chrysomeloidea). Pakaluk J Slipinski SA Eds Biol. Phylogeny Classif. Coleopt. Pap. Celebr. 80th Birthd. Roy CrowsonMuzeum Inst. Zool. PAN Warszawa 2: 559–631.

Robert, C. A., N. Veyrat, G. Glauser, G. Marti, G. R. Doyen et al., 2012 A specialist root herbivore exploits defensive metabolites to locate nutritious tissues: a root herbivore exploits plant defences. Ecol. Lett. 15: 55–64.

Rostant, W. G., N. Wedell, and D. J. Hosken, 2012 Transposable elements and insecticide resistance, pp. 169–201 in Advances in Genetics, Elsevier.

Russell, S. J., and J. LaMarre, 2018 Transposons and the PIWI pathway: genome defense in gametes and embryos. Reproduction 156: R111–R124.

Saha, A., J. A. Mitchell, Y. Nishida, J. E. Hildreth, J. A. Ariberre et al., 2015 A trans-Dominant Form of Gag Restricts Ty1 Retrotransposition and Mediates Copy Number Control. J. Virol. 89: 3922–3938.

Santiago, N., C. Herráiz, J. R. Goñi, X. Messeguer, and J. M. Casacuberta, 2002 Genome-wide analysis of the *Emigrant* family of MITEs of *Arabidopsis thaliana*. Mol. Biol. Evol. 19: 2285–2293.

Sappington, T. W., B. D. Siegfried, and T. Guillemaud, 2006 Coordinated *Diabrotica* genetics research: accelerating progress on an urgent insect pest problem. Am. Entomol. 52: 90–97.

Sasai, H., M. Ishida, K. Murakami, N. Tadokoro, A. Ishihara et al., 2009 Species-Specific Glucosylation of DIMBOA in Larvae of the Rice Armyworm. Biosci. Biotechnol. Biochem. 73: 1333–1338.

Schoville, S. D., Y. H. Chen, M. N. Andersson, J. B. Benoit, A. Bhandari et al., 2018 A model species for agricultural pest genomics: the genome of the Colorado potato beetle, *Leptinotarsa decemlineata* (Coleoptera: Chrysomelidae). Sci. Rep. 8: 1931.

Smit, A. F. A., R. Hubley, and P. Green, 1996 Repeat-Masker open-3.0. http://www.repeatmasker.org.

Souza, D., J. A. Peterson, R. J. Wright, and L. J. Meinke, 2020 Field efficacy of soil insecticides on pyrethroid-resistant western corn rootworms (*Diabrotica virgifera virgifera* LeConte). Pest Manag. Sci. 76: 827–833.

Souza, D., B. C. Vieira, B. K. Fritz, W. C. Hoffmann, J. A. Peterson et al., 2019 Western corn rootworm pyrethroid resistance confirmed by aerial application simulations of commercial insecticides. Sci. Rep. 9: 6713.

Talla, V., A. Suh, F. Kalsoom, V. Dincă, R. Vila et al., 2017 Rapid increase in genome size as a consequence of transposable element hyperactivity in wood-white (*Leptidea*) butterflies. Genome Biol. Evol. 9: 2491–2505.

Thomas-Bulle, C., M. Piednoël, T. Donnart, J. Filée, D. Jollivet et al., 2018 Mollusc genomes reveal variability in patterns of LTR-retrotransposons dynamics. BMC Genomics 19: 821.

Toepfer, S., T. Haye, M. Erlandson, M. Goettel, J. G. Lundgren et al., 2009 A review of the natural enemies of beetles in the subtribe Diabroticina (Coleoptera: Chrysomelidae): implications for sustainable pest management. Biocontrol Sci. Technol. 19: 1–65.

Tomaszewska, P., T. K. Pellny, L. M. Hernández, R. A. C. Mitchell, V. Castiblanco et al., 2021 Flow cytometry-based determination of ploidy from dried leaf specimens in genomically complex collections of the tropical forage grass *Urochloa* s. l. Genes 12: 957

Voronova, A., V. Belevich, A. Jansons, and D. Rungis, 2014 Stress-induced transcriptional activation of retrotransposon-like sequences in the Scots pine (*Pinus sylvestris L*.) genome. Tree Genet. Genomes 10: 937–951.

Wang, X., X. Fang, P. Yang, X. Jiang, F. Jiang et al., 2014 The locust genome provides insight into swarm formation and long-distance flight. Nat. Commun. 5: 2957.

Wang, S., M. D. Lorenzen, R. W. Beeman, and S. J. Brown, 2008 Analysis of repetitive DNA distribution patterns in the *Tribolium castaneum* genome. Genome Biol. 9: R61.

Wang, L., J. Wang, Y. Ma, P. Wan, K. Liu et al., 2019 Transposon insertion causes cadherin mis-splicing and confers resistance to Bt cotton in pink bollworm from China. Sci. Rep. 9: 7479.

Wang, X., Y. Xu, S. Zhang, L. Cao, Y. Huang et al., 2017 Genomic analyses of primitive, wild and cultivated citrus provide insights into asexual reproduction. Nat. Genet. 49: 765–772.

Weinstock, G. M., G. E. Robinson, R. A. Gibbs, G. M. Weinstock, G. M. Weinstock et al., 2006 Insights into social insects from the genome of the honeybee *Apis mellifera*. Nature 443: 931–949.

Wendel, J. F., and S. R. Wessler, 2000 Retrotransposon-mediated genome evolution on a local ecological scale. Proc. Natl. Acad. Sci. 97: 6250–6252.

Wicker, T., F. Sabot, A. Hua-Van, J. L. Bennetzen, P. Capy et al., 2007 A unified classification system for eukaryotic transposable elements. Nat. Rev. Genet. 8: 973–982.

Yan, H., A. Bombarely, and S. Li, 2020 DeepTE: a computational method for de novo classification of transposons with convolutional neural network. Bioinformatics 36: 4269–4275.

Zhang, S., S. Gu, X. Ni, and X. Li, 2019 Genome size reversely correlates with host plant range in *Helicoverpa* species. Front. Physiol. 10: 29.

